# Does wearable neurofeedback training improve memory aptitudes in healthy adults? ––A systematic review and meta-analysis

**DOI:** 10.1101/2025.11.03.685283

**Authors:** Makoto Hagihara, Reiji Ohkuma, Tatsuro Fujimaki, Masako Tamaki, Mitsuaki Takemi, Maro G. Machizawa

**Affiliations:** Department of Physiological Sciences, School of Life Science, The Graduate University for Advanced Studies (SOKENDAI), Aichi, Japan; Division of Neural Dynamics, Department of System Neuroscience, National Institute for Physiological Sciences, Aichi, Japan; Graduate School of Human Sciences, Waseda University, Saitama, Japan; Graduate School of Science and Technology, Keio University, Kanagawa, Japan; Cognitive Somnology RIKEN Hakubi Research Team, RIKEN Cluster for Pioneering Research, Saitama, Japan; Graduate School of Advanced Science and Engineering, Hiroshima University, Hiroshima, Japan; Center for Brain, Mind and KANSEI Sciences Research, Hiroshima University, Hiroshima, Japan; Xiberlinc Inc., Tokyo, Japan

**Keywords:** Neurofeedback training, Memory, Healthy adults, Electroencephalogram (EEG), Functional near-infrared spectroscopy (fNIRS), Meta-analysis, Randomized controlled trials

## Abstract

Neurofeedback training (NFT) provides individuals with real-time feedback on their brain activity, enabling voluntary modulation. Interest in NFT for memory enhancement has surpassed the cumulative evidence in healthy adults, especially for modalities feasible outside of laboratory settings. To address this gap, this systematic review and meta-analysis focused on electroencephalography (EEG) and functional near-infrared spectroscopy (fNIRS), both of which are portable and widely accessible. Randomized controlled trials published between January 1990 and September 2022 were retrieved from PubMed, Scopus, Web of Science, and APA PsycInfo. Of the 44 eligible studies, 24 reported sufficient data for inclusion in the meta-analysis. Using a random-effects model, we found that NFT produced a small-to-moderate improvement in memory performance (standardized mean difference = 0.28; 95% confidence interval = 0.065–0.48; *t*(11.9) = 2.86; *p* = 0.014), with moderate heterogeneity. No publication bias was detected. Subgroup meta-analyses revealed significant effects in younger adults, for EEG-based protocols, and in tasks assessing verbal and short-term memory tasks. Meta-regression identified a positive association between daily session duration and effect size, with sessions longer than 30 minutes more likely to produce positive effects. As effects also varied by targeted brain regions and EEG frequency bands used for neurofeedback, protocol differences likely contributed to the observed variability in outcomes. Long-term follow-up is warranted to determine the persistence of NFT effects.

## 1. Introduction

Memory plays a critical role in human cognition and complex behavior. It is essential for acquiring and retaining new information, making decisions informed by past experiences, and executing daily tasks. As such, improving memory capacity can have significant benefits in areas such as academic performance, workplace productivity, and personal efficiency (<u>Cowan, 2014</u>).

The concept of "memory" has been defined in various ways by researchers (<u>Baddeley, 2000</u>; <u>Tulving, 1983</u>). It is a multifaceted and complex process comprising multiple subsystems, and it can be categorized by retention duration into short-term memory (STM), long-term memory (LTM), working memory, and episodic memory. Each subsystem contributes to different cognitive tasks and behaviors. For example, STM is involved in the immediate retention of information, while episodic memory handles the recall of personal experiences (<u>Squire, 2004</u>). Despite the diversity among memory types, many enhancement efforts have focused on techniques that modulate the brain’s neural activity.

One such technique is neurofeedback training (NFT). NFT is a method that enables individuals to self-regulate their brain activity by receiving real-time feedback on specific neural patterns and is commonly implemented with noninvasive sensors. This feedback is typically delivered through visual, auditory, or tactile cues, and it is based on monitoring brain activity via electroencephalography (EEG), functional near-infrared spectroscopy (fNIRS), or other neuroimaging techniques. Although much of the research on NFT has focused on improving attention and executive function (<u>Egner & Gruzelier, 2004</u>; <u>Gruzelier, 2014</u>), recent studies suggest that it may also enhance working and episodic memory by modulating oscillatory rhythms implicated in memory processing and encoding, such as theta (4–8 Hz) and alpha (8–12 Hz) rhythms (<u>Klimesch, 1999</u>; <u>Sitaram et al., 2017</u>; <u>Escolano et al., 2011</u>). These observations motivate a systematic evaluation of NFT for memory enhancement.

Despite these promising results, it remains uncertain whether the benefits and practical effectiveness of NFT extend to healthy adult populations. Most previous research has targeted clinical cohorts (<u>Kober et al., 2015</u>a; <u>Trambaiolli et al., 2021</u>; <u>Lavy et al., 2021</u>) or reported broad cognitive outcomes without differentiating memory subtypes (<u>Lee et al., 2013</u>; <u>Yeo et al., 2018</u>), leaving the specific effects of NFT in the context of memory systems in healthy individuals less well characterized. Furthermore, the factors contributing to NFT-related improvements in memory performance remain unclear. To address these gaps, a systematic synthesis examining how methodological conditions such as frequency band, target brain region, and training duration relate to variability in NFT efficacy across memory subtypes is warranted.

Here we conducted a systematic review of the effects of NFT on memory aptitudes in healthy adults, focusing on EEG- and fNIRS-based protocols because these wearable modalities are portable, widely available, and increasingly accessible outside laboratory settings. This focus enables evaluation of NFT approaches that are most feasible for practical use in everyday environments and consumer applications. We also performed a meta-analysis to quantify the overall effect and to examine effects across memory subtypes (e.g., STM, LTM, verbal, visuospatial), NFT protocols (e.g., increasing alpha or beta power), and control conditions. This work differs from five recent systematic reviews that addressed the effects of NFT on memory aptitudes in healthy adults (<u>Yeh et al., 2021</u>; <u>Yeh et al., 2022</u>; <u>Viviani and</u> <u>Vallesi, 2021</u>; <u>Matsuzaki et al., 2023</u>; <u>Jackson et al., 2023</u>) in several respects. Prior meta-analyses focused on specific frequency bands such as alpha (<u>Yeh et al., 2021</u>) and theta (<u>Yeh et al., 2022</u>) or on single memory domains such as working memory (<u>Viviani and Vallesi, 2021</u>; <u>Matsuzaki et al., 2023</u>) or episodic memory (<u>Jackson et al., 2023</u>). Consequently, prior reviews did not evaluate NFT’s effects across multiple memory subtypes within a unified framework, nor did they comprehensively compare protocol features in healthy adults. Moreover, the comparative effectiveness of NFT relative to control conditions, such as sham NFT or alternative cognitive interventions, remains insufficiently explored. Understanding how different NFT protocols affect various aspects of memory and how these effects compare to other interventions is crucial for determining the broader applicability of NFT in enhancing cognitive performance.

## 2. Methods

This systematic review and meta-analysis adhered to Chapter 4 of the Minds Manual for Clinical Practice Guideline Development (<u>Minds Manual Developing Committee ed., 2021</u>). The study was preregistered with the International Prospective Register of Systematic Reviews (PROSPERO: CRD42022381936; <u>Machizawa et al., 2022</u>). As this is a review study, it did not require approval from an ethics committee or informed consent from participants.

### 2.1 Database search

Three of the six authors (M.H., R.O., T.F., Ma.T., Mi.T., and M.M.) independently searched for English-language articles investigating NFT effects on memory aptitudes using four electronic databases (PubMed [R.O. and T.F.], Scopus [M.H. and R.O.], Web of Science [M.H. and T.F.], and APA PsycInfo [R.O. and T.F.]). All database searches were conducted in September 2022.

Search strings combined memory-related and NFT-related terms. Memory-related terms included “memor*” OR “remember*” OR “recall*” OR “reminiscenc*” OR “retent*” OR “recollect*” OR “recognition” OR “learning”. In contrast, NFT-related terms were “neuro*feedback” OR “neuropsychological feedback” OR “neuronal feedback” OR “neural feedback” OR “bio*feedback” OR “brain machine interface” OR “brain computer interface” OR “BMI” OR “BCI”. To eliminate irrelevant studies, we excluded any articles in which the following keywords appeared in the title or keywords: "animal*" OR "rodent*" OR "mouse" OR "mice" OR "monkey*" OR "rat" OR "macaque*" OR "marmoset*" OR "child*" OR "kid*" OR "infant*" OR "EMG" OR "motor*" OR "sport*" OR "athletic*" OR "Deja Vu" OR "priming*" OR "attention*" OR ("pattern recognition" AND "machine learning") OR "sensory memory" OR "emotion* recognition" OR "body mass index". Supplementary Table 1 details the exact search queries used for each database.

Although our preregistration (<u>Machizawa et al., 2022</u>) included "sensory memory" as one of the target domains, it was later excluded during the review process. This decision was based on the recognition that sensory memory, by definition, reflects ultra-short-term information retention and is not typically regarded as a modifiable component of memory function (<u>Vallet & Desrichard, 2016</u>). This review aimed to evaluate the efficacy of NFT on memory systems with behavioral relevance, such as STM and LTM. Accordingly, "sensory memory" was excluded from the final search terms and analysis to ensure conceptual consistency with the primary research question.

### 2.2 Screening procedures

The screening process occurred in two phases. In the first phase, studies were excluded based on their title and abstract. In the second phase, full texts were reviewed for eligibility. During full-text screening, the following data were extracted and considered for eligibility: study design, participant health, intervention type, control condition, and outcome measures related to memory aptitudes.

Eligibility criteria were as follows: (1) Population: healthy adults aged 18 years or older; (2) Intervention: NFT using noninvasive wearable brain imaging technique (i.e., EEG or fNIRS) aimed at enhancing memory aptitudes; (3) Comparison: any; (4) Outcome: memory aptitude assessed via validated measures; and (5) Article type: original studies published in peer-reviewed English-language journals. Three authors (M.H., R.O., and T.F.) independently conducted the screening process. Articles were divided into three subsets, each reviewed by two of the three authors (M.H. and R.O., R.O. and T.F., or T.F. and M.H.). In case of disagreement between these two authors, resolution was sought through review by the remaining co-authors (M.H., R.O., T.F., or M.M., depending on the pair). Eligible studies were excluded from the meta-analysis if (1) effect size data for experimental and control groups were unavailable; (2) the study design was not a randomized controlled trial (RCT); or (3) the study was assessed as having a high risk of bias or substantial indirectness (see Sections 2.4 & 2.5 for criteria).

### 2.3 Data extraction

Three authors (M.H., R.O., and T.F.) extracted data from each eligible study for both qualitative synthesis and meta-analysis. Extracted variables included participant demographics, intervention type, control condition, memory outcome measures, and any reported adverse events. Adverse events were defined as any unfavorable medical occurrences during the study period, including symptoms such as headache, nausea, or dizziness (<u>Higgins et al., 2019</u>). Memory outcomes were categorized into four domains: (1) STM (e.g., digit span task): measuring the capacity to temporarily retain and manipulate a sequence of numbers presented in a specific order (<u>Nouchi et al., 2022</u>); (2) LTM (e.g., free recall task): assessing the ability to retrieve previously learned information after a delay without cues (<u>Rozengurt et al., 2017</u>); (3) Visuospatial memory (e.g., delayed matching to sample task): measuring the ability to remember and identify a visual or spatial pattern after a delay by matching it to a sample from a set of options (<u>Campos Da Paz et al.,</u> <u>2018</u>); and (4) Verbal memory (e.g., word-pair test): measuring the ability to learn, store, and recall pairs of words after a delay, assessing associative memory for verbal information (<u>Wei et al., 2017</u>). Three authors (M.H., R.O., and T.F.) independently classified the memory domains. Disagreements were resolved through discussion with additional authors (Ma.T. and M.M.) until consensus was reached.

For meta-analysis, we extracted outcome measures of memory aptitudes after the last NFT session. Here, we used changes in scores from pre- to post-NFT when available. If change scores were unavailable, we used post-intervention scores. Each experiment was treated as an independent trial. For each trial, we extracted means, standard deviations, and sample sizes from the experimental and control groups. From either these values or *t*-values, the standardized mean difference (SMD) was calculated using Hedges’ *g*. When relevant data were unavailable, we contacted corresponding authors to request the missing information. To ensure consistency in effect direction, we reversed SMD signs when lower scores reflected better performance (e.g., reaction time), so that higher SMD values always indicated superior memory outcomes in the NFT group compared to controls.

### 2.4 Risk of bias assessment

Two authors (M.H. and R.O.) independently assessed the risk of bias in all studies for which SMDs were available, using the Cochrane Collaboration tool (<u>Higgins et al., 2011</u>). This tool categorizes risk of bias as high, unclear, or low across six domains: selection bias, performance bias, detection bias, attrition bias, reporting bias, and others (e.g., conflict of interest, early termination of trials without reasonable reasons, incorrect sample size estimation, or flawed statistical methods). We assigned two points for high risk, one for unclear risk, and zero for low risk. An overall score ≥8 or ≥2 domains rated high was categorized as high risk; scores ≤4 with ≤1 high-risk domains were categorized as low risk; all other cases were categorized as moderate risk. Only studies with low or moderate risk were included in the meta-analysis. A risk-of-bias summary table was generated using “*robvis*” (<u>McGuinness & Higgins, 2021</u>).

### 2.5 Indirectness assessment

We evaluated indirectness across four domains: participant selection, intervention, control conditions, and relevance of the targeted outcomes. Each domain was rated as low, high, or unclear based on predefined criteria. For participant selection, indirectness was rated as low when studies provided sufficient detail on sample characteristics and included healthy adult participants; as high when selection involved stratification based on specific traits (e.g., IQ or cognitive test scores) or conditions (e.g., prior sleep deprivation); and as unclear when the selection process lacked adequate description. For the intervention domain, indirectness was rated as low when studies implemented NFT in isolation; as high when interventions incorporated additional components beyond NFT (e.g., post-training sleep periods that could confound outcomes); and as unclear when protocols lacked sufficient detail or could not be independently verified. For control conditions, we rated indirectness as low when studies employed active controls such as general memory training or sham neurofeedback; as high when comparators involved no-intervention conditions, or when order effects could not be ruled out; and as unclear when comparator details were insufficient. For the relevance of the targeted outcomes, we rated indirectness as low when studies used established quantitative measures directly linked to memory aptitude; as high when outcomes were assessed using subjective measures or questionnaires with fewer than five response levels; and as unclear when outcome measurement procedures were insufficiently described. All discrepancies in rating were resolved through discussion until consensus was reached.

### 2.6 Meta-analysis

We conducted a random-effects pairwise meta-analysis, pooling results from all trials that reported comparable outcome measures. Statistical heterogeneity was assessed using Cochran’s Q and the *I*^2^ statistic. Publication bias was assessed both visually using a funnel plot and statistically using Begg’s and Egger’s tests, with a significance threshold of *p* < 0.1, in accordance with the Minds Manual for Clinical Practice Guideline Development (<u>Minds Manual Developing Committee ed., 2021</u>).

Subgroup analyses were performed to evaluate the effects of NFT based on memory subtype (STM, LTM, visuospatial memory, or verbal memory), age group (18–60 years or >60 years), recording modality (EEG or fNIRS), measured brain region (frontal, central, parietal, or occipital), target frequency band (theta, alpha, or beta), and control condition (sham NFT or general training). Subgroup analysis was not conducted for gamma-band NFT, as only one study met the inclusion criteria for the meta-analysis. Because several studies reported multiple outcomes from the same participant samples, we applied a cluster-robust variance estimation method (<u>Fisher & Tipton, 2015</u>) using the *clubSandwich* package in R. This method accounts for the statistical dependency of multiple effect sizes within individual studies.

A meta-regression analysis was conducted to assess the dose-response relationship between daily NFT duration and effect size. If the NFT duration was reported as a range, we used the midpoint (e.g., 35 minutes for 30–40 minutes). We also examined the relationship between effect size and the timing of post-intervention memory assessments. All meta-analyses were performed using R version 4.1.2, with the following packages: “*metafor*” ver. 3.4.0 (<u>Viechtbauer, 2010</u>), “*meta*” ver. 5.5.0 (<u>Balduzzi et al., 2019</u>), and “*clubSandwich*” ver. 0.5.8 (<u>Fisher & Tipton, 2015</u>).

### 2.7 Multiple correspondence analysis

In addition to category-specific subgroup meta-analyses, we conducted a multiple correspondence analysis (MCA) to identify latent relationships among cross-category study features and their association with intervention effectiveness. MCA is a nonparametric multivariate technique for exploring associations among categorical variables (<u>Lorenzo-Seva & van de Velden, 2019</u>). Because few trials used fNIRS or targeted the gamma band, the MCA was restricted to EEG-based studies targeting theta, alpha, or beta bands to ensure sufficient sample size. The MCA included the following six categorical variables: namely age (younger or older), memory category (visuospatial or verbal), memory duration (STM or LTM), EEG frequency band (theta, alpha, or beta), targeted brain region (frontal, central, parietal, or occipital), and a binary effectiveness classification (effective or null). As for the effectiveness, each SMD value was classified as ‘effective’ if the lower bound of the 95% confidence interval (CI) for the SMD exceeded 0.

Otherwise, it was classified as ‘null’. MCA was performed using the “*MultipleCar*” toolbox (<u>Lorenzo-Seva</u> <u>& van de Velden, 2019</u>) run on MATLAB 2024a (Mathworks, Natick, MA). We interpreted the factor map by examining proximities among category points and orientation relative to the Effective–Null axis.

## 3. Results

### 3.1 Search results

Figure 1 depicts the study selection process. We identified 3,927 articles from four databases (PubMed, Scopus, Web of Science, and APA PsycInfo) using predefined search strings. Of these, 3,839 records were excluded based on their titles or abstracts. The remaining 88 articles were then screened for eligibility by full-text assessment. Forty-four articles were excluded at this stage for the following reasons: absence of memory-related outcome measures (*n* = 11; including studies that used only mental rotation tasks, which we classified as spatial visualization rather than memory), lack of a control group (*n* = 11), ineligible article types (*n* = 9), no implementation of neurofeedback (*n* = 8), or inclusion of clinical populations (*n* = 5). Consequently, 44 articles met the inclusion criteria and were retained for qualitative synthesis.

**Figure 1.**
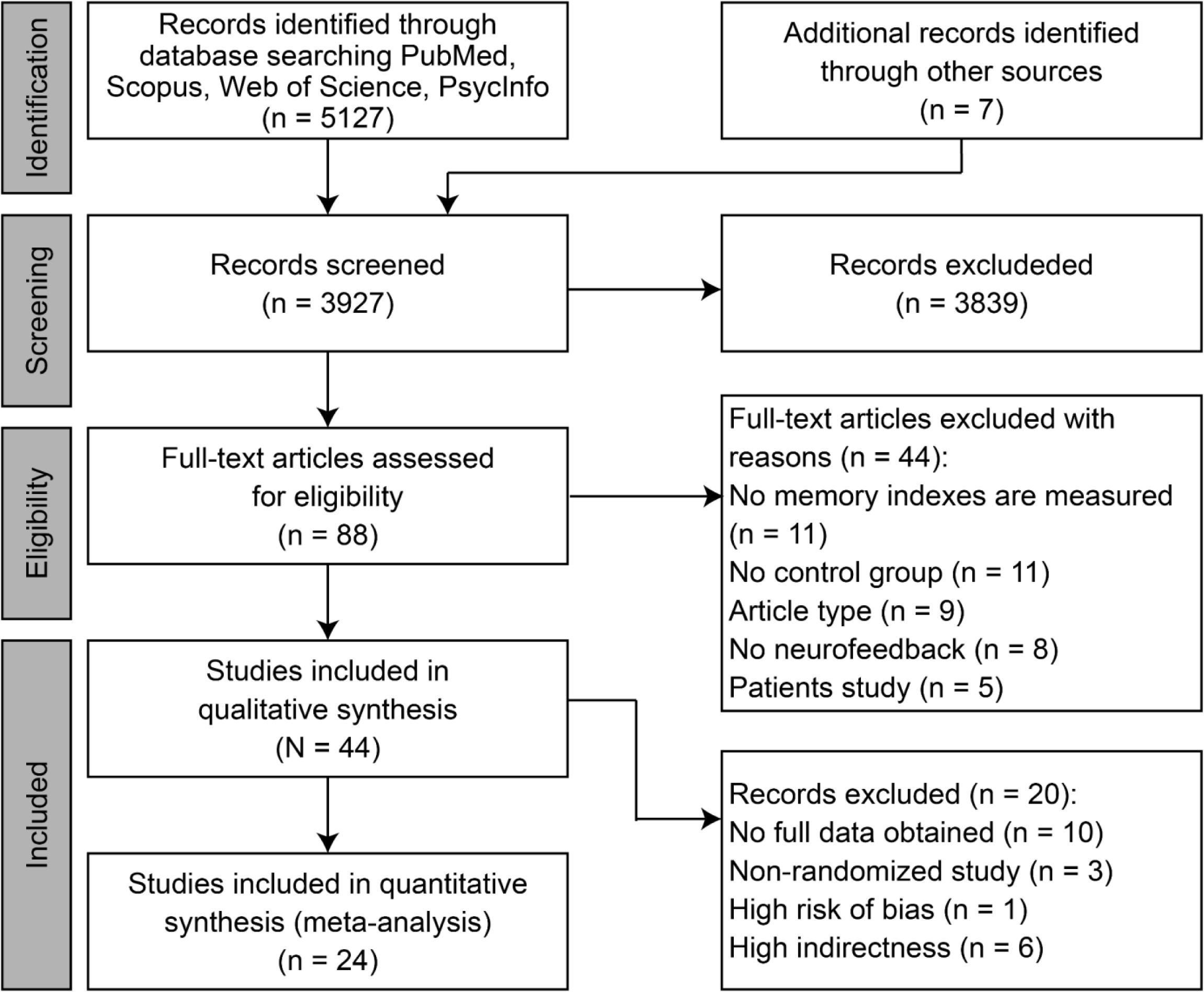
PRISMA flowchart illustrating the study selection process.

### 3.2 Study and sample characteristics

The basic characteristics of the studies included in the qualitative synthesis are summarized in Table 1. All 44 articles were published between 1975 and 2022, with more than half published after 2013 (Fig. 2A). Thirty-seven studies employed EEG-based neurofeedback, while seven used fNIRS (Fig. 2B). Regarding the memory domain, 27 studies targeted STM alone, four examined LTM alone, and 13 addressed both STM and LTM (Fig. 2C).

**Figure 2.**
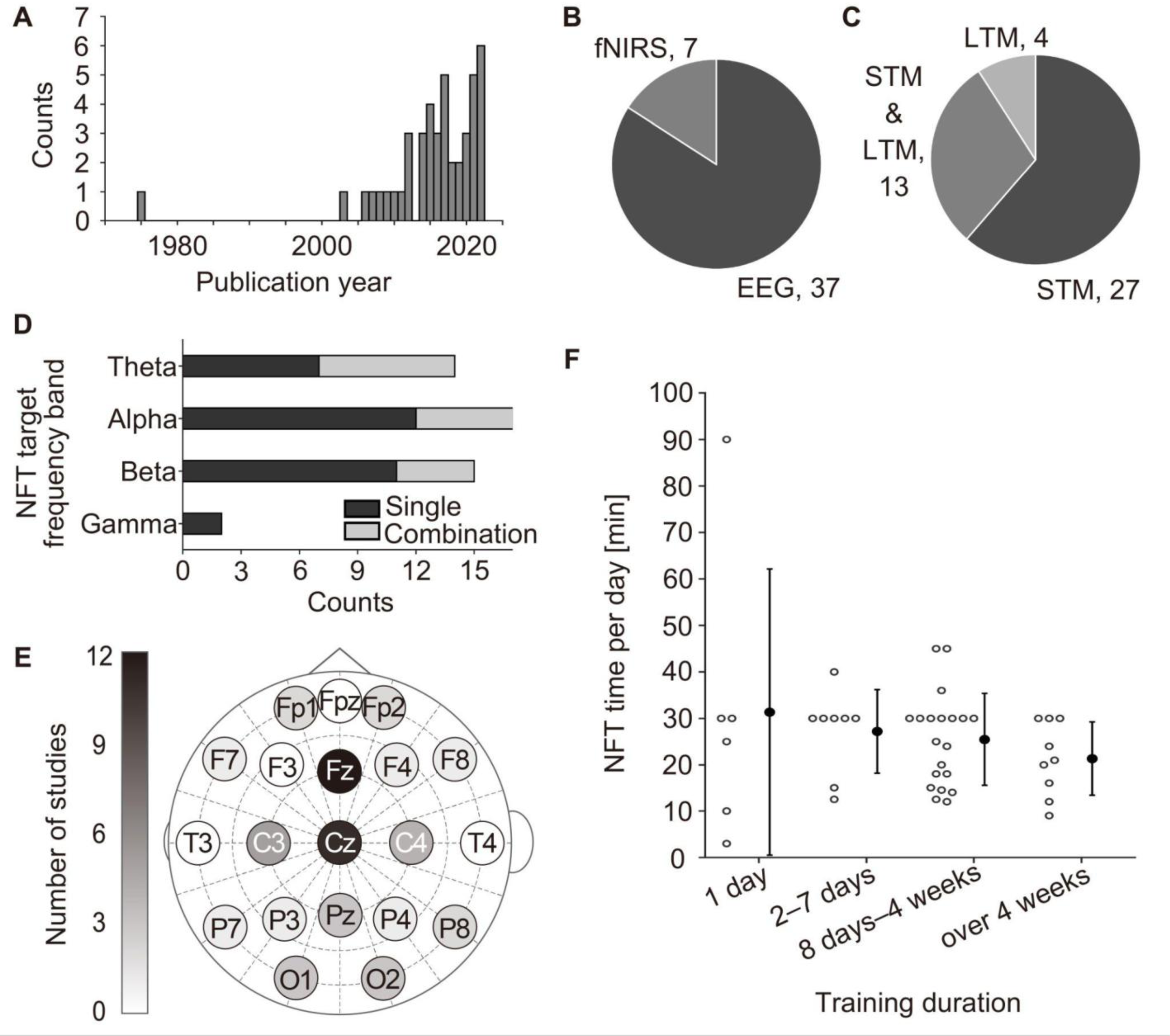
Overview of the studies included in the qualitative synthesis. (A) Publication year distribution of the included studies. (B) Neuroimaging modalities used for neurofeedback training (NFT). EEG: electroencephalography; fNIRS: functional near-infrared spectroscopy. (C) Memory subtypes assessed (short-term memory (STM), long-term memory (LTM), or both). (D) Frequency bands targeted in EEG-based NFT, distinguishing single vs. combined band protocols. (E) Scalp locations used as NFT targets in EEG studies. Channels used as sham or nonactive conditions were excluded. (F) Daily NFT session durations plotted against the total intervention length. Open circles indicate individual studies; black dots and error bars represent the group means and standard deviations, respectively.

**Table 1.**
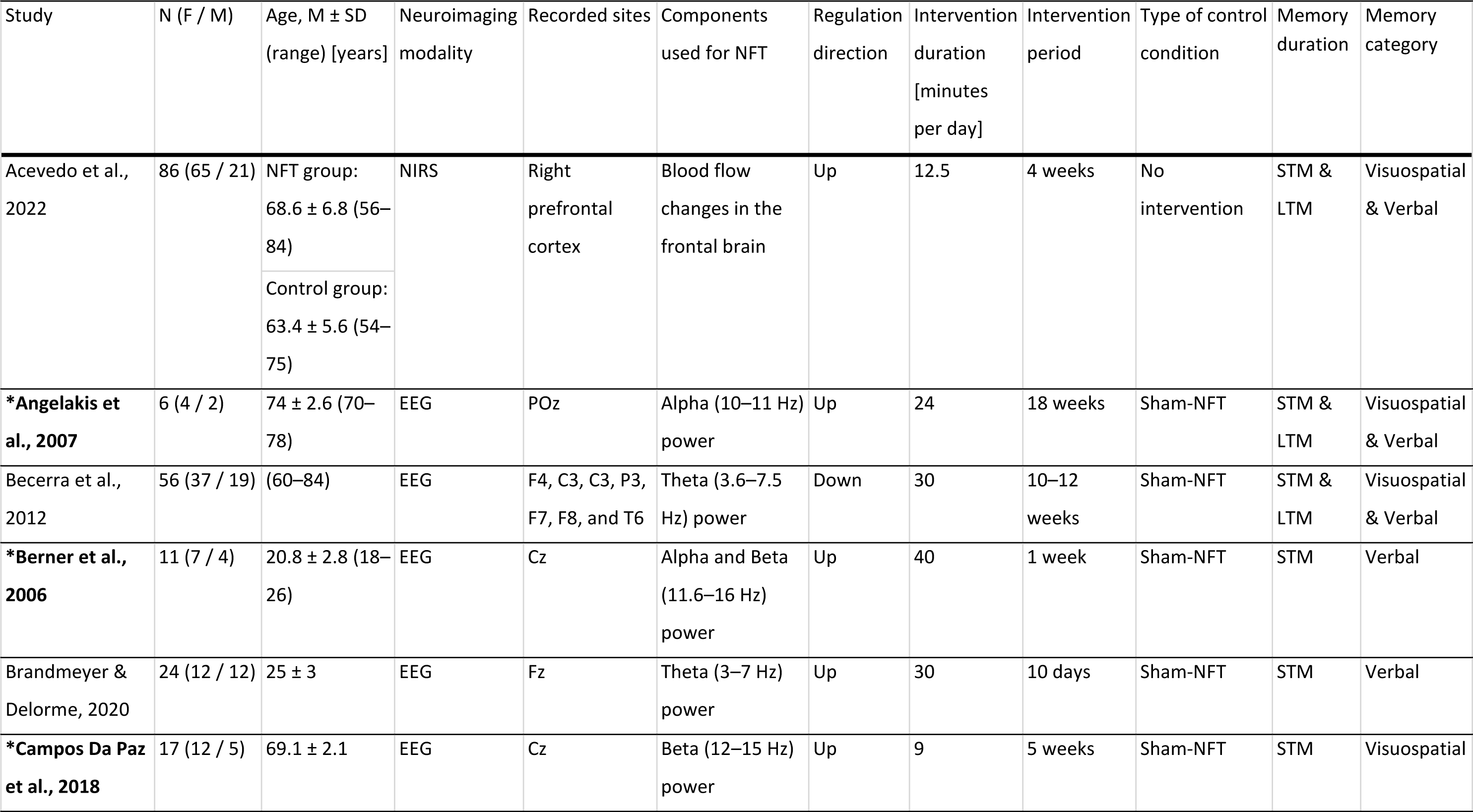
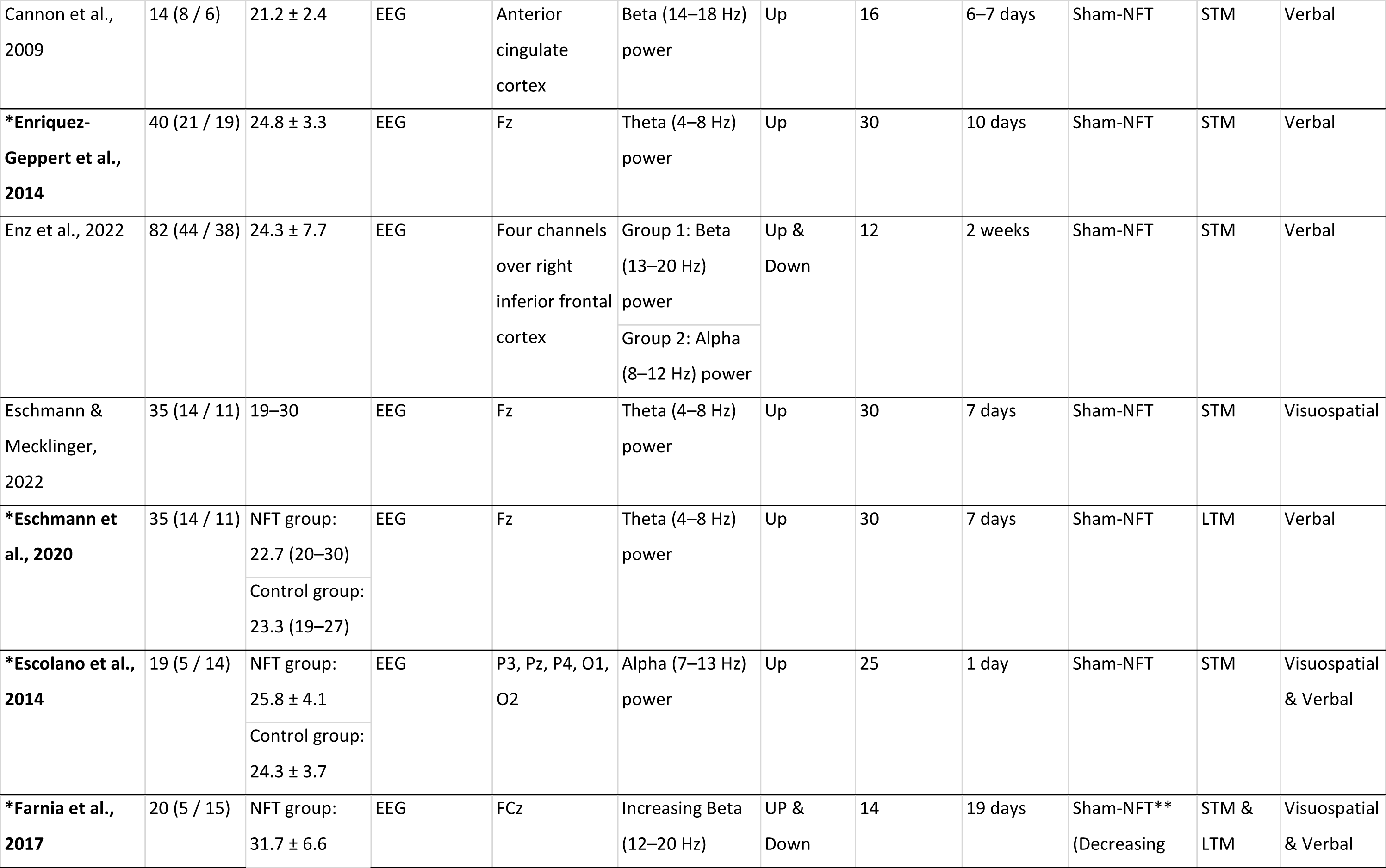
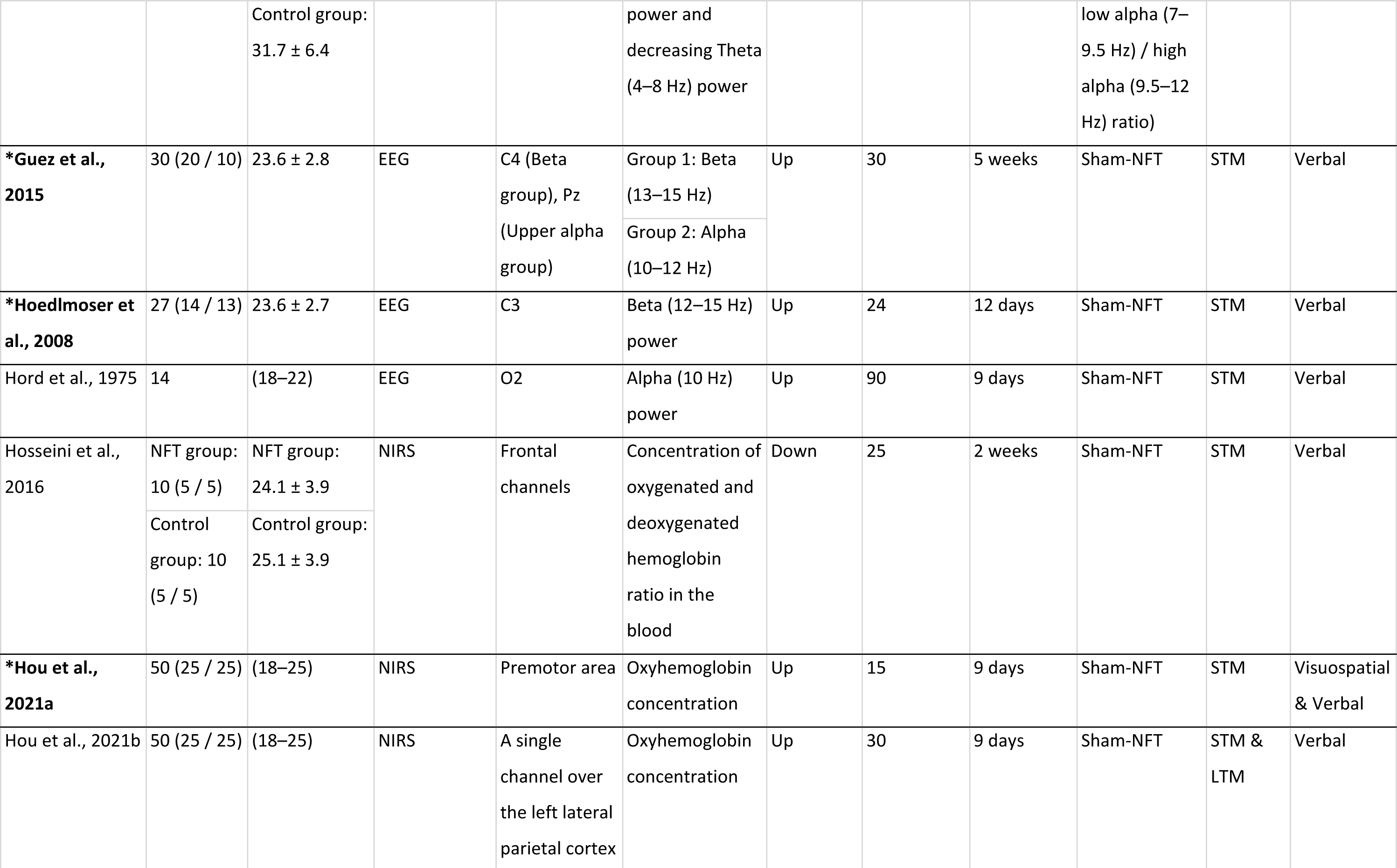
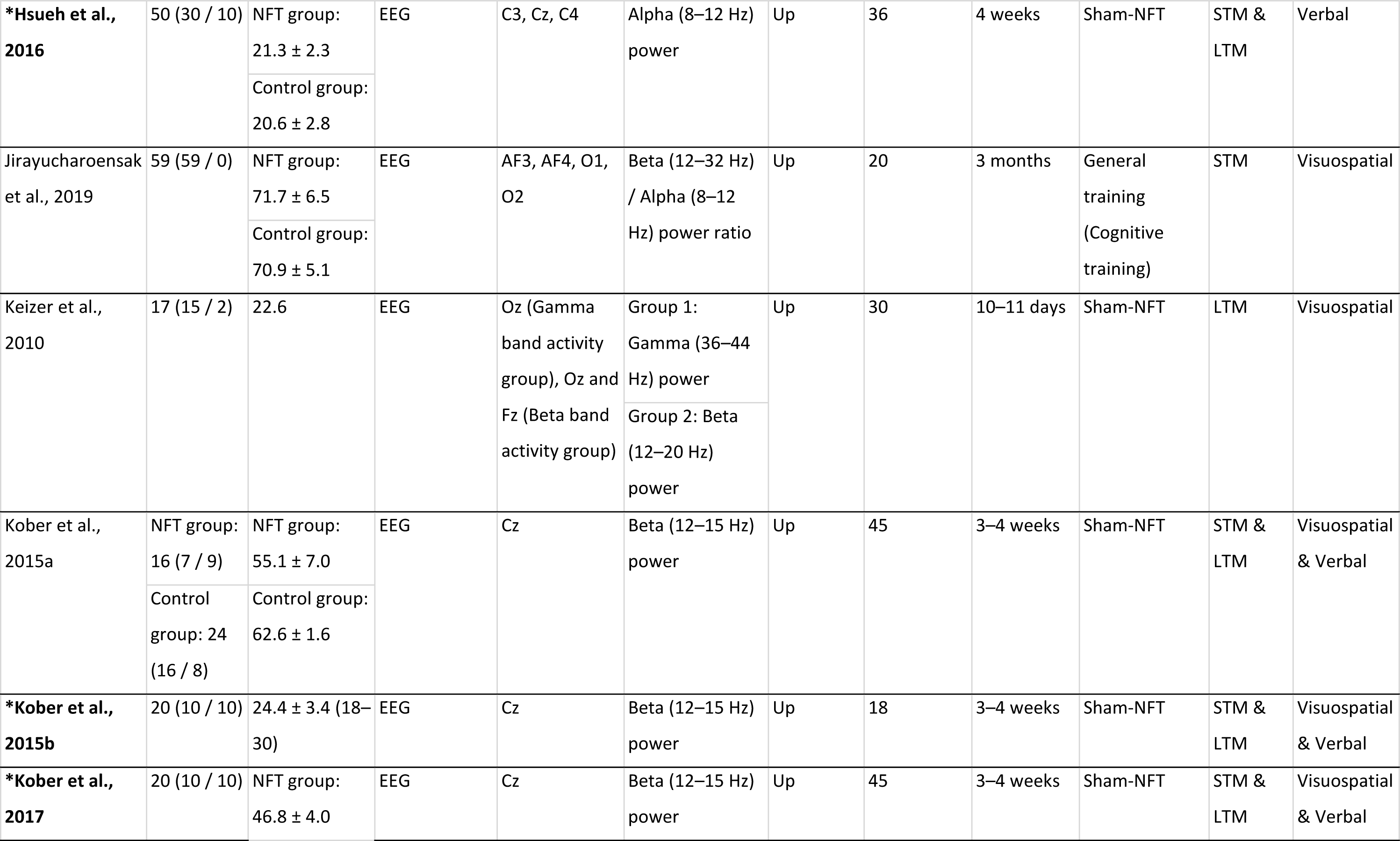
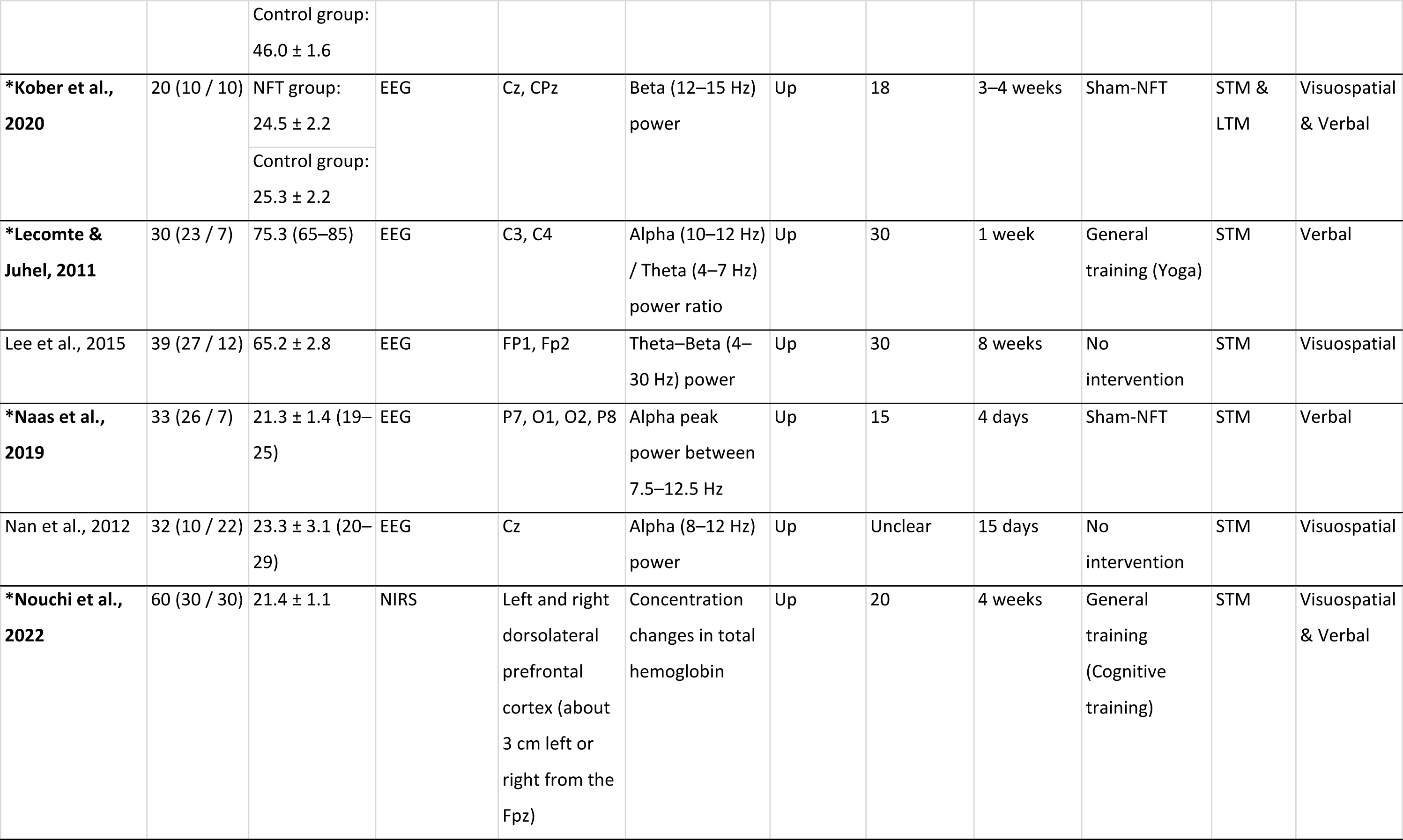
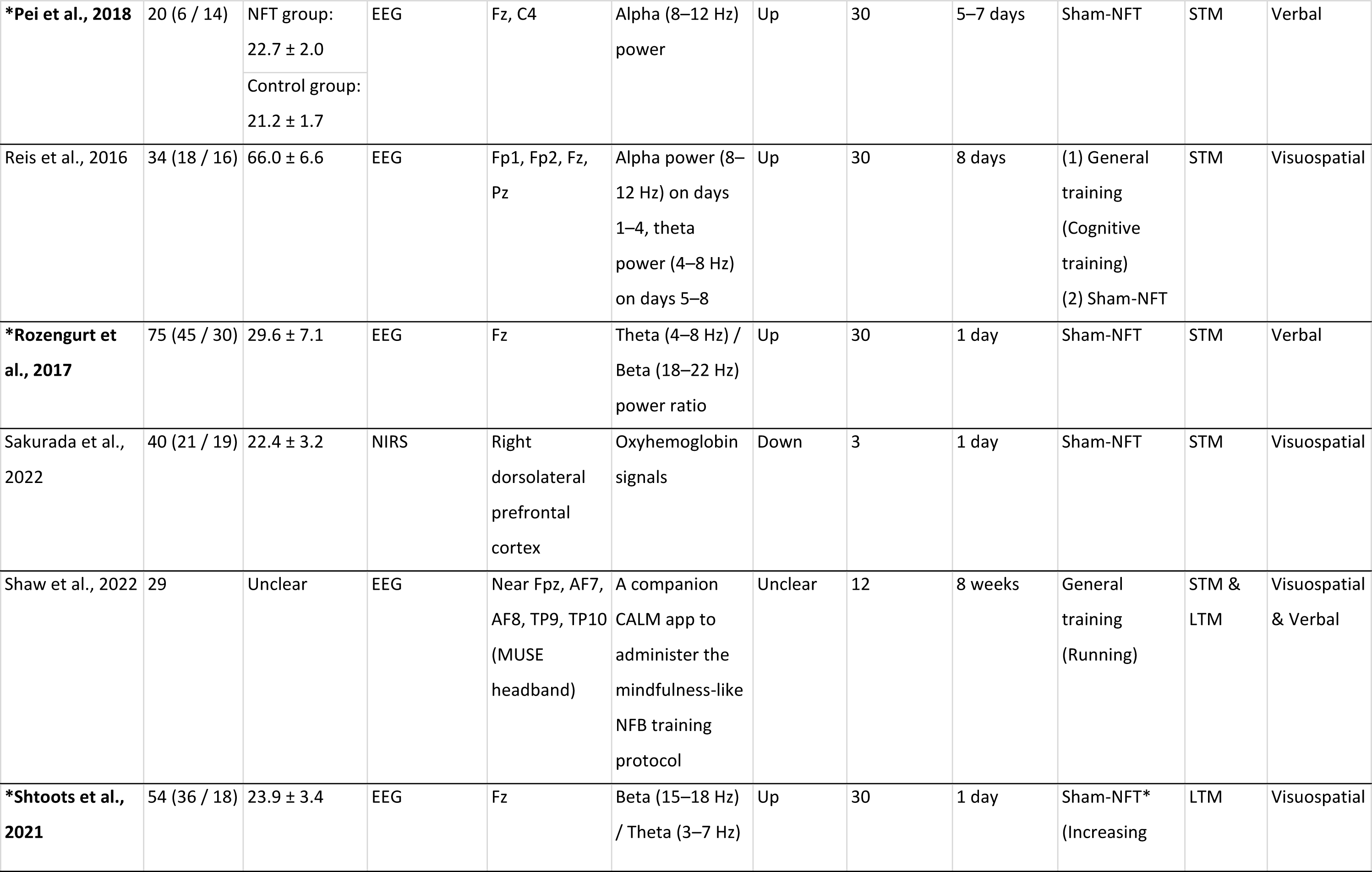
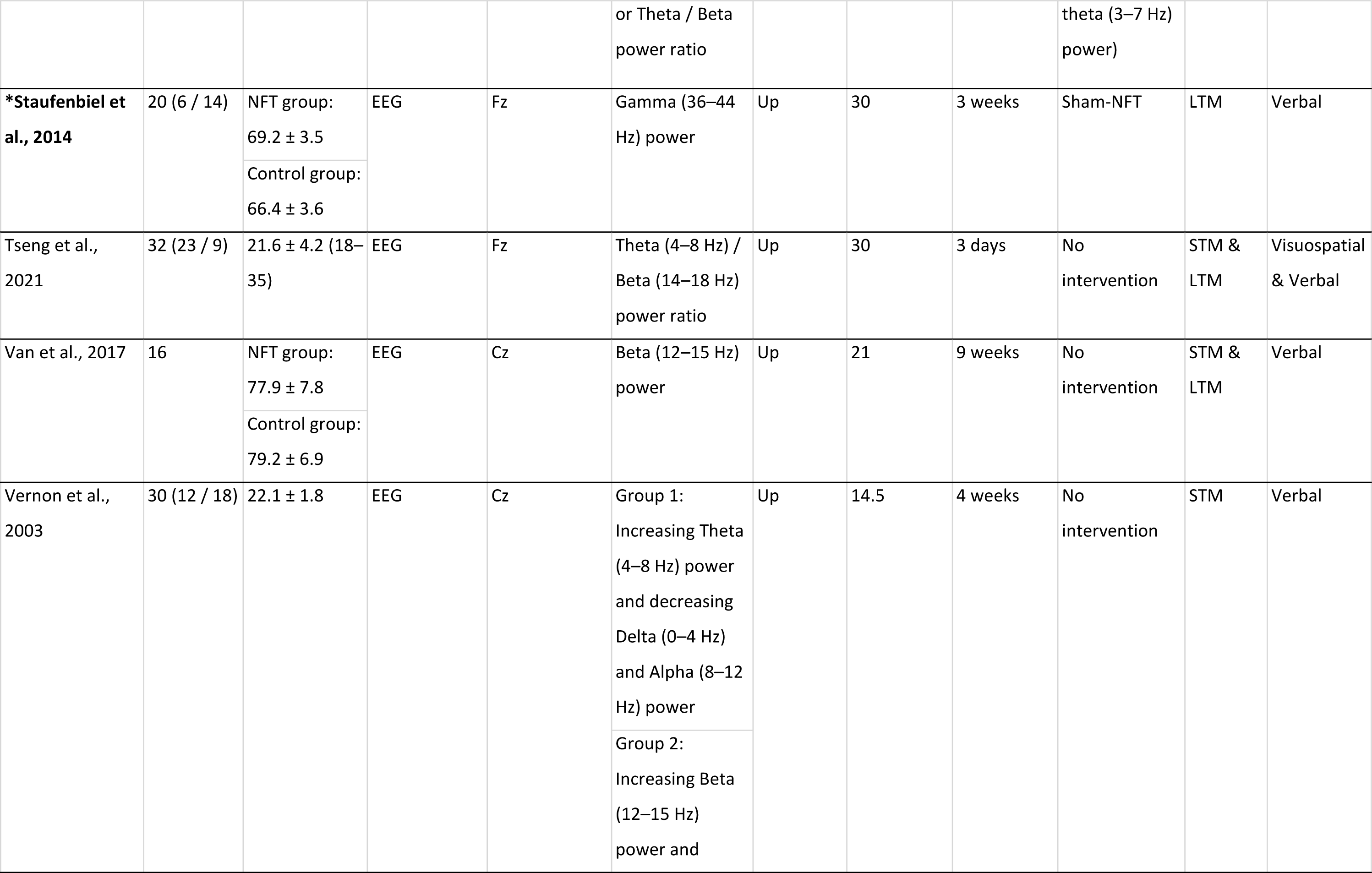
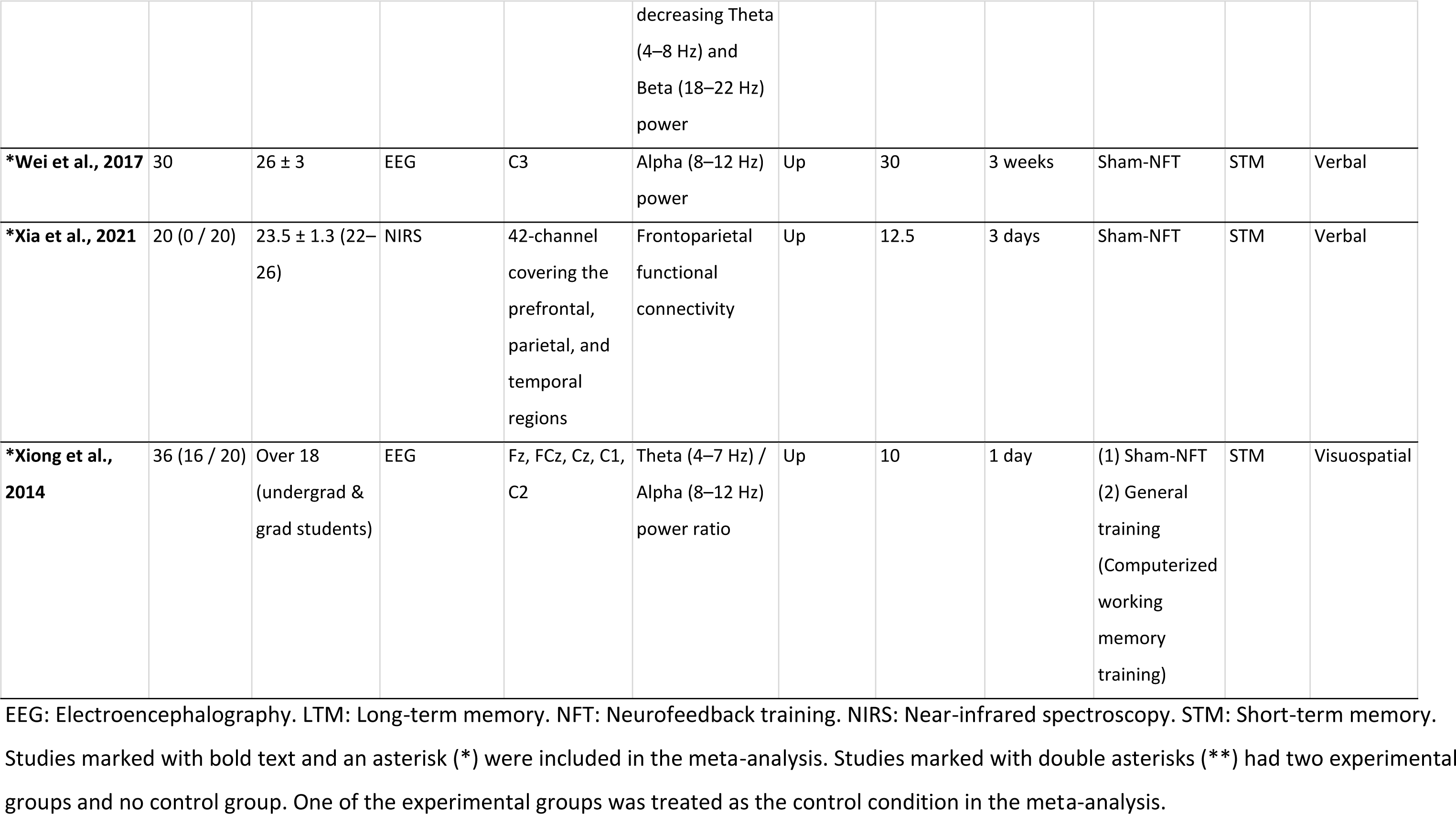
Experimental characteristics of the studies included in the qualitative synthesis.

Neurofeedback protocols in EEG-based studies predominantly targeted modulation of theta, alpha, or beta frequency bands. Gamma-band neurofeedback was rare, with only 2 of 44 studies aiming to modulate gamma activity. Most studies focused on modulating a single frequency band rather than multiple bands. Specifically, among the studies targeting alpha (*n* = 17) or beta (*n* = 18) activity, 10 studies in each group used a single-band protocol (Fig. 2D). For the theta band, single-frequency targeting was also more common (5 of 15 studies), although a subset employed combined-frequency protocols.

Electrode placements in EEG-based NFT studies were concentrated around the midline frontal region, with Fz being the most frequently used target. Additional commonly targeted electrode sites included the vertex (Cz), central (C3, C4), occipital (O1, O2), midline parietal (Pz), frontopolar (Fp1, Fp2), occipitotemporal (T5, T6), parietal (P3, P4), right frontal (F4), and inferior frontal (F7, F8) regions (Fig. 2E). Intervention durations spanned from one day to 18 weeks, with daily sessions ranging from 3 to 90 minutes (Fig. 2F). Regarding study design, 34 studies used sham NFT as a control, 6 employed no-intervention controls, and 6 included general cognitive training as the comparator.

### 3.3 Adverse events

Among the 44 studies included in the qualitative synthesis, only four explicitly reported on the presence or absence of adverse events. None of these studies reported any adverse effects occurring during or following NFT.

### 3.4 Risk of bias

From the 44 studies in the qualitative synthesis, we evaluated the risk of bias in 31 studies. These 31 studies met two inclusion criteria: they were randomized controlled trials (RCTs) and reported sufficient data to compute effect sizes. Thus, they were considered eligible for meta-analysis, contingent upon subsequent risk of bias and indirectness assessment. The remaining 13 studies were excluded from risk of bias evaluation because 10 lacked sufficient data for effect size estimation, and 3 were not randomized trials.

Among the 31 studies evaluated, the overall risk of bias was high in one study, moderate in 11 studies, and low in 18 studies. Only three studies explicitly described the method of random sequence generation and were rated as low risk for selection bias (Fig. 3, column A). The remaining studies lacked sufficient detail on randomization procedures and were thus classified as unclear risk. Regarding allocation concealment (Fig. 3, column B), only one study described its method clearly and was rated low risk; all others were judged unclear. For performance bias (Fig. 3, column C), 11 studies were single-blinded or unblinded and consequently rated high risk. Seven studies were double-blinded and classified as low risk, while 12 studies did not describe their blinding procedures and were rated as unclear. Detection bias (Fig. 3, column D) was mostly unclear, as most studies did not report whether outcome assessors were blinded. One study was explicitly unblinded. Attrition bias was generally low across studies (Fig. 3, column E): 28 were rated low risk, one was unclear, and one was judged to have violated attrition criteria. Reporting bias (Fig. 3, column F) was predominantly unclear; 29 studies lacked preregistration, while one was rated low risk for adhering to a prespecified protocol. Regarding other biases (Fig. 3, column G), three studies were rated as low risk based on sufficient detail regarding potential conflicts of interest, sample size estimation, early trial termination, or statistical integrity. The remaining 27 studies were rated as unclear due to insufficient information.

**Figure 3.**
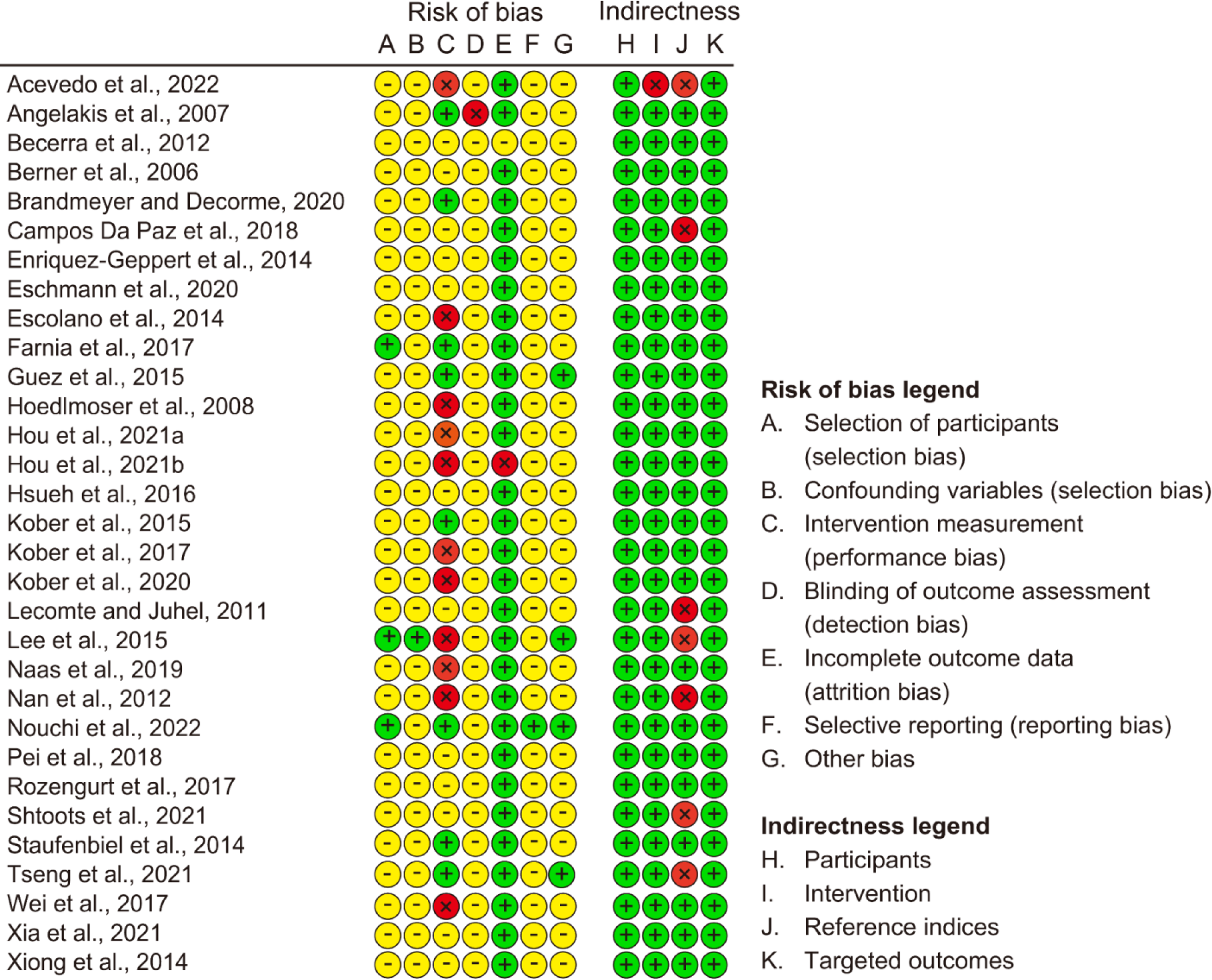
Risk of bias and indirectness assessment results. Columns A–G denote bias domains, and H–K indicate indirectness components. Green plus (+) indicates low risk/little indirectness; yellow minus (–) indicates unclear; red cross (×) indicates high risk/substantial indirectness.

### 3.5 Indirectness assessment

After excluding one study due to a high risk of bias, indirectness was evaluated across the remaining 30 studies eligible for meta-analysis. These assessments are visualized in Figure 3 (columns H–K). All 30 studies were rated as low for participant selection, as they provided sufficient detail about sample characteristics. Regarding the intervention, 29 studies were rated as low, while one study was rated as high due to the combination of NFT with general memory training, which precluded isolation of the specific effects of NFT. For control conditions, 24 studies were rated as low risk, whereas six were rated high risk because they used passive or no-intervention controls, potentially compromising comparability. All studies were considered low risk regarding the relevance of the targeted outcomes. Based on these criteria, six studies were excluded from the meta-analysis for exhibiting substantial indirectness in one or more domains.

### 3.6 Overall effects of NFT on memory performance

A total of 73 RCTs from 24 studies (649 participants) were included in the meta-analysis using a random-effects model. Overall, NFT yielded a small but statistically significant improvement in memory performance (Fig. 4A; SMD = 0.28, 95% CI = 0.065–0.48, *t* (11.9) = 2.86, *p* = 0.014). No evidence of publication bias was identified, based on visual inspection of funnel plots (Supplementary Fig. 1), Begg’s test (*τ* = 0.10, *p* = 0.20), and Egger’s test (*Z* = -0.020, *p* = 0.98). Heterogeneity across studies was moderate (*Q* (72) = 119.78, *p* = 0.003, *I*² = 40.0%).

**Figure 4.**
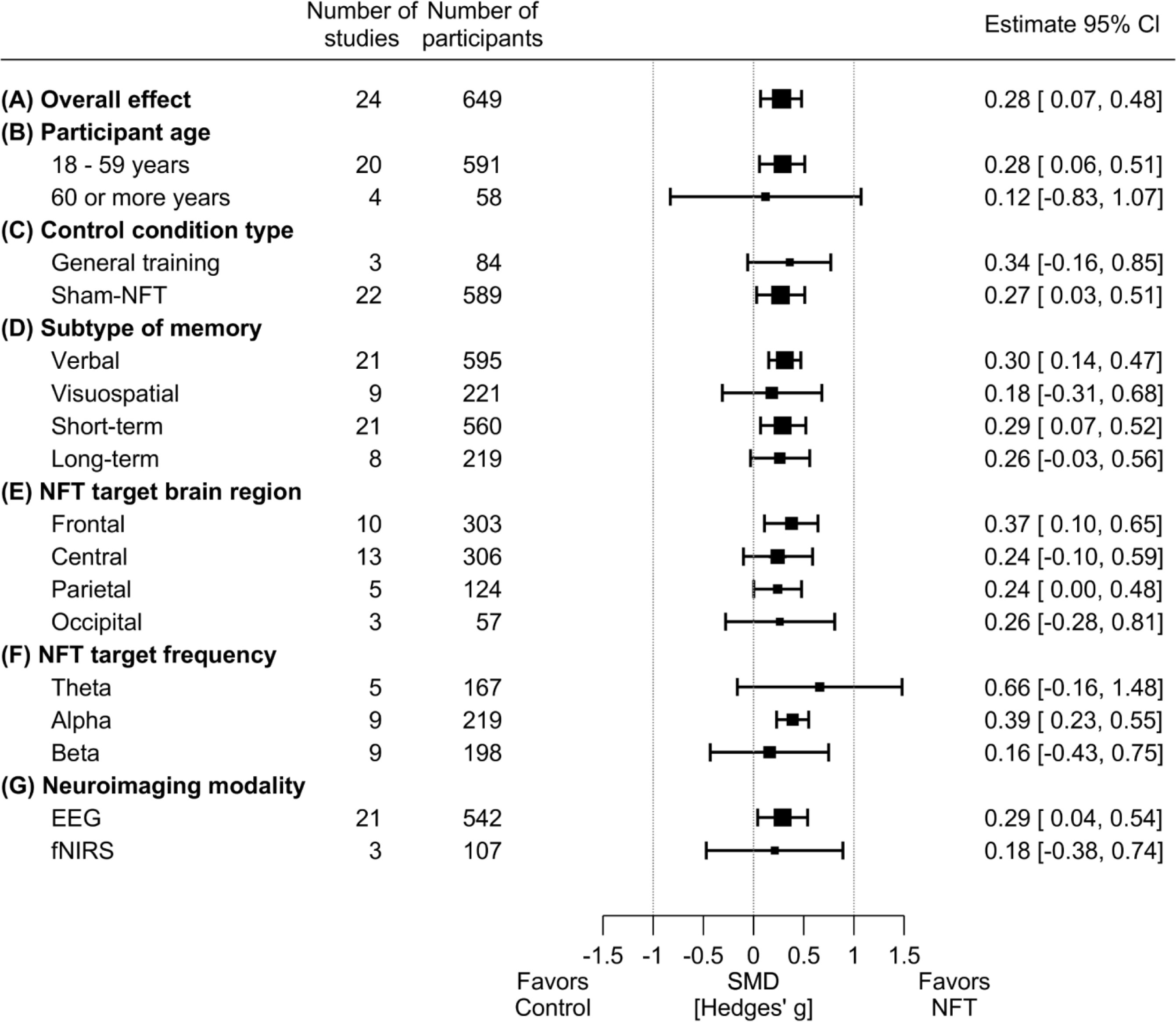
Forest plots of the meta-analysis examining the effect of NFT on memory performance (Hedges’ g) across the overall sample and the prespecified subgroups. (A) Overall effect. (B) Participants’ age. (C) Control condition. (D) Memory subtype. (E) Targeted brain region. (F) EEG frequency band (no studies targeted delta; one targeted gamma). (G) Neuroimaging modalities. Black squares represent pooled effect size estimates, with square size proportional to the number of contributing studies; horizontal lines show 95% confidence intervals (CIs). Positive values favor NFT over the control. SMD, standardized mean difference.

### 3.7 Subgroup meta-analyses

Subgroup meta-analyses were conducted to examine the influence of participant age, control type, memory subtype, target brain region, frequency band, and neuroimaging modality.

#### 3.7.1 Participant age (Fig. 4B)

The studies were stratified into two age groups: younger adults (18–59 years) and older adults (≥60 years). No statistically significant differences in NFT effect sizes were observed between these subgroups. In younger adults, NFT demonstrated a small but statistically significant effect (20 studies; SMD = 0.29, 95% CI = 0.020–0.56, *t* (10.6) = 2.75, *p* = 0.020), with moderate heterogeneity (*Q* (66) = 116.9, *p* = 0.0001, *I*² = 43.6%). In contrast, no significant effect was detected in older adults (4 studies; SMD = 0.12, 95% CI = - 0.83–1.072, *t* (1.6) = 0.69, *p* = 0.57), with no detectable heterogeneity (*Q* (5) = 2.21, *p* = 0.82, *I*² = 0%).

#### 3.7.2 Control condition type (Fig. 4C)

NFT effects were compared across two control conditions: general cognitive training and sham neurofeedback. No statistically significant difference in effect sizes was observed between these subgroups. Relative to general training, NFT yielded a non-significant effect (3 studies; SMD = 0.34, 95% CI = -0.16–0.85, *t* (1.5) = 4.25, *p* = 0.086), with substantial heterogeneity (*Q* (6) = 18.1, *p* = 0.0060, *I*² = 68.2%). Compared to sham NFT, a small but significant effect was observed (22 studies; SMD = 0.27, 95% CI = 0.027–0.51, *t* (10.4) = 2.47, *p* = 0.032), with moderate heterogeneity (*Q* (65) = 101.5, *p* = 0.0025, *I*² = 36.2%).

#### 3.7.3 Memory subtype (Fig. 4D)

Memory outcomes were categorized as verbal, visuospatial, STM, or LTM. No statistically significant differences were detected between subtypes. NFT produced moderate effects on verbal memory (21 studies; SMD = 0.30, 95% CI = 0.14–0.47, *t* (13.2) = 3.95, *p* = 0.0016) with moderate heterogeneity (*Q* (52) = 95.4, *p* = 0.0002, *I*² = 45.8%) and STM (21 studies; SMD = 0.29, 95% CI = 0.071–0.52, *t* (13.4) = 2.84, *p* = 0.014) with moderate heterogeneity (*Q* (52) = 105.2, *p* < 0.0001, *I*² = 51.3%). No statistically significant effects were observed for visuospatial memory (9 studies; SMD = 0.18, 95% CI = -0.31–0.68, *t* (5.1) = 0.95, *p* = 0.39; heterogeneity: *Q* (18) = 23.4, *p* = 0.18, *I*² = 15.1%) or LTM (8 studies; SMD = 0.26, 95% CI = -0.034–0.559, *t* (4.8) = 2.30, *p* = 0.072; heterogeneity: *Q* (20) = 15.1, *p* = 0.77, *I*² = 0%).

#### 3.7.4 Target brain region (Fig. 4E)

NFT target sites were grouped into four regions: frontal, central, parietal, and occipital. NFT targeting the frontal cortex showed a moderate and significant effect (10 studies; SMD = 0.38, 95% CI = 0.10–0.65, *t* (6.7) = 3.21, *p* = 0.016) with high heterogeneity (*Q* (20) = 59.5, *p* < 0.0001, *I*² = 67.3%). A small but statistically significant effect was observed for parietal targeting (5 studies; SMD = 0.24, 95% CI = 0.0024– 0.48, *t* (2.6) = 3.48, *p* = 0.049) with no heterogeneity (heterogeneity: *Q* (8) = 7.00, *p* = 0.54, *I*² = 0%). No significant effects were found for central targeting (13 studies; SMD = 0.24, 95% CI = -0.10–0.59, *t* (6.0) = 1.73, *p* = 0.13; heterogeneity: *Q* (49) = 72.3, *p* = 0.017, *I*² = 31.5%) or occipital targeting (3 studies; SMD = 0.26, 95% CI = -0.28–0.81, *t* (1.2) = 4.46, *p* = 0.11; heterogeneity: *Q* (4) = 1.42, *p* = 0.84, *I*² = 0%).

#### 3.7.5 Target frequency (Fig. 4F)

EEG-based NFT studies were categorized by target frequency: theta, alpha, or beta. NFT targeting the alpha band demonstrated a small but statistically significant effect (9 studies; SMD = 0.39, 95% CI = 0.23– 0.55, *t* (4.9) = 6.19, *p* = 0.0017), with low heterogeneity (*Q* (16) = 20.9, *p* = 0.18, *I*² = 19.9%). No significant effects were observed for beta-band NFT (9 studies; SMD = 0.16, 95% CI = -0.43–0.75, *t* (3.4) = 0.80, *p* = 0.48; heterogeneity: *Q* (37) = 37.1, *p* = 0.47, *I*² = 0%) or theta-band NFT (5 studies; SMD = 0.66, 95% CI = - 0.16–1.48, *t* (2.8) = 2.66, *p* = 0.082; heterogeneity: *Q* (8) = 33.2, *p* < 0.0001, *I*² = 77.5%).

#### 3.7.6 Neuroimaging modality (Fig. 4G)

NFT studies were categorized as EEG-based or fNIRS-based. A small but statistically significant effect was detected for EEG-based NFT (21 studies; SMD = 0.29, 95% CI = 0.039–0.54, *t* (9.68) = 2.59, *p* = 0.028), accompanied by moderate heterogeneity (*Q* (64) = 104.9, *p* = 0.0010, *I*² = 39.4%). By contrast, fNIRS-based NFT did not produce a statistically significant effect (3 studies; SMD = 0.18, 95% CI = -0.38–0.74, *t* (1.67) = 1.66, *p* = 0.26), although heterogeneity was moderate (*Q* (7) = 14.1, *p* = 0.049, *I*² = 49.9%).

### 3.8 Meta-regression analysis

Meta-regression revealed a statistically significant positive association between effect size and daily NFT session duration (24 studies; *β* = 0.016, 95% CI = 0.0001–0.031, *t* (3.6) = 2.95, *p* = 0.049; Fig. 5A). Importantly, effect sizes exceeding 0.2, a commonly used benchmark for a small but meaningful effect in behavioral sciences (<u>Cohen, 1988</u>), were observed only when session durations surpassed 30 minutes, suggesting a possible trend toward greater efficacy with longer training durations. By contrast, no significant association was observed between effect size and the timing of post-training memory assessments (15 studies; *β* = 0.009, 95% CI = -0.14–0.15, *t* (1.6) = 0.32, *p* = 0.78; Fig. 5B).

**Figure 5.**
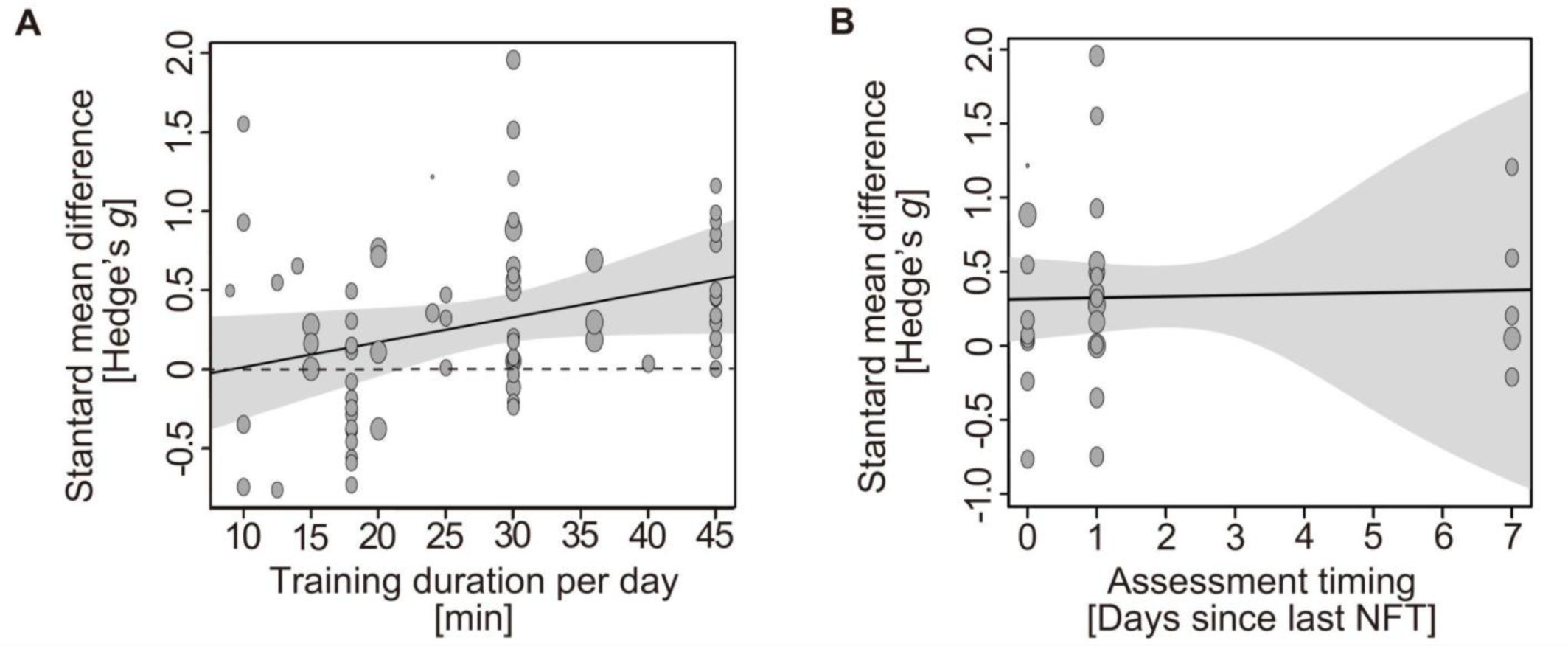
Meta-regression analyses of NFT effectiveness. (A) SMD (Hedges’ g) as a function of daily NFT session duration, showing a significant positive association (*p* = 0.049). (B) SMD as a function of the delay (in days) between NFT completion and memory testing timing, showing no significant association. The shaded areas represent the 95% CIs of the fitted trend lines.

### 3.9 Multiple correspondence analysis

Figure 6 presents the multiple correspondence analysis (MCA) map, in which Dimensions 1 and 2 accounted for 22.0% and 19.4% of the total variance, respectively. The NFT-effectiveness axis (Effective vs. Null) aligned closely with Dimension 1, such that categories positioned on the positive side (to the right of the origin) tended to co-occur with larger observed effects. Verbal outcomes and STM were positioned nearer the Effective category point than visuospatial outcomes and LTM. Younger adults were also closer to the Effective category point than older adults. These patterns were consistent with the subgroup meta-analysis, supporting the convergent validity of the MCA. Along Dimension 2, targeted brain regions were segregated so that frontal and central clustered together on one side, opposite parietal and occipital. The map also hinted at characteristic pairings between anatomical regions and frequency bands—specifically frontal-theta, central-beta, and parietal-alpha.

**Figure 6.**
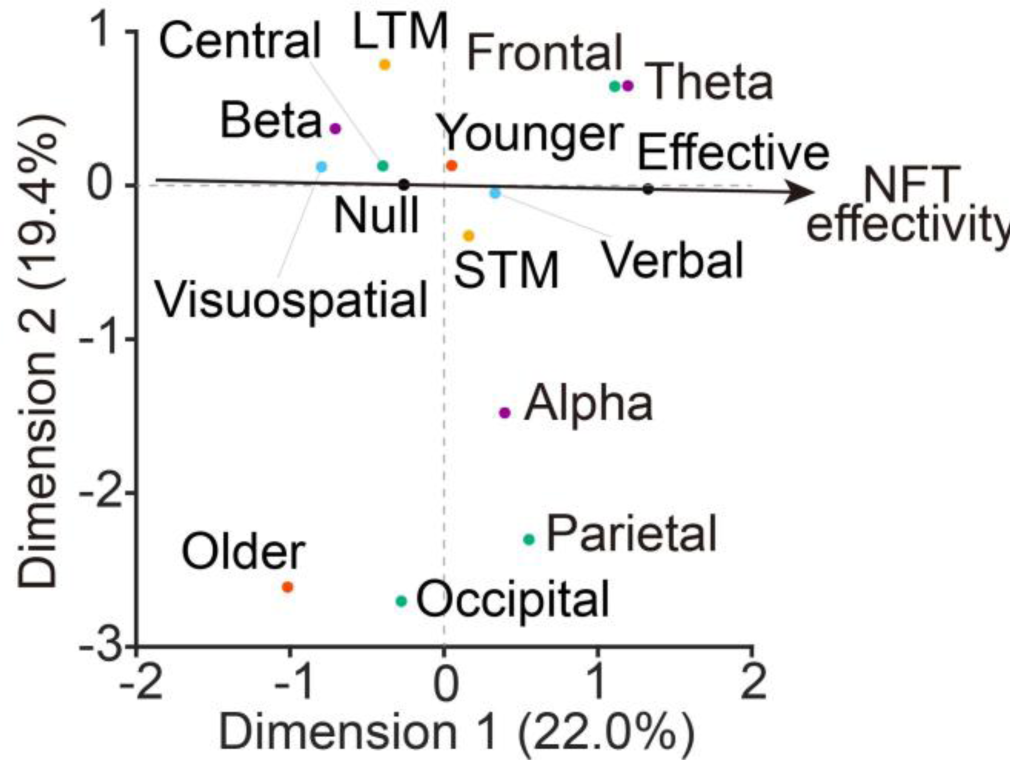
Multiple correspondence analysis (MCA) map of categorical study features in EEG-based NFT studies. The categories included effectiveness (*Effective* or *Null*; black), age (younger or older adults; blue), memory category (verbal or visuospatial; purple), memory duration (short-term [STM] or long-term [LTM]; yellow), EEG frequency band (theta, alpha, or beta; magenta), and channel region (frontal, central, parietal, or occipital; green). Effectiveness was defined as *Effective* when the lower bound of the 95% CI for the SMD exceeded 0; otherwise it was defined as *Null*. The black arrow indicates the Effective–Null axis. The distances between category points reflect associations; closer points indicate stronger co-occurrence, whereas larger separations indicate weaker associations. Dimensions 1 and 2 explained 22.0% and 19.4% of the variance, respectively.

## 4. Discussion

This systematic review and meta-analysis provide quantitative support for the efficacy of NFT in enhancing memory functions among healthy adults. An initial literature search identified 3927 articles, of which 44 studies met the criteria for qualitative synthesis. Across 24 studies included in the final meta-analysis, comprising 73 RCTs, NFT was associated with a small but statistically significant improvement in memory performance. Serious adverse events were not reported in the four studies that monitored them. Taken together, the evidence suggests that NFT may offer modest cognitive benefits under specific conditions, with a favorable safety profile that warrants further investigation in both research and applied contexts.

### 4.1 Age-based differences in NFT efficacy

The subgroup meta-analysis revealed that NFT did not yield statistically significant effects in older adults, whereas significant improvements were observed in younger adults. This discrepancy may be attributable to limited sample sizes in older adults, as well as greater heterogeneity in cognitive status among aging populations. Notably, behavioral research has shown that older adults retain the capacity to improve multitasking performance through structured cognitive training (<u>Anguera et al., 2013</u>). Therefore, the absence of a pooled effect here does not preclude benefits in this age group. Instead, the pattern suggests that age-appropriate NFT targets and dosing may be necessary. Future randomized trials with sufficient sample sizes should test this hypothesis. From a neurobiological perspective, younger adults generally exhibit higher baseline levels of neuroplasticity (<u>Pauwels et al., 2018</u>; <u>Tymofiyeva & Gaschler, 2021</u>), which may enhance their responsiveness to NFT protocols. This neuroplastic advantage may partially explain the more pronounced effects observed in younger cohorts.

### 4.2 Comparison with general training and sham NFT

In direct comparisons, NFT did not outperform general cognitive training. By contrast, NFT showed a small but statistically significant advantage over sham neurofeedback. These contrasts may reflect limited statistical power and precision due to small sample sizes within certain subgroups. Collectively, the findings suggest that NFT produces effects beyond placebo but does not clearly outperform generalized cognitive training in improving memory outcomes.

### 4.3 Differences by memory type, target brain region, and EEG frequency band

Subgroup analyses revealed differential effects based on memory type, targeted brain region, and EEG frequency band. Specifically, the memory-type subgroup showed moderate effects for STM and no significant effects for LTM. This dissociation likely reflects the greater susceptibility of short-timescale processes to NFT than LTM. Because EEG-based NFT has been reported to enhance attentional functions (<u>Kimura et al., 2024</u>), it is reasonable to expect corresponding benefits for STM, which depends heavily on prefrontal attentional control. In contrast, LTM depends on distributed encoding, consolidation, and retrieval processes mediated by hippocampal-cortical interactions, which may attenuate the effects of short-duration training and contribute to greater inter-individual variability (<u>Sitaram et al., 2017</u>). Moreover, verbal memory outcomes showed significant improvement following NFT, whereas visuospatial outcomes did not, and effect size estimates were highly heterogeneous across studies. Among the nine visuospatial studies, only two reported statistically significant improvements, one targeting central beta (<u>Kober et al., 2017</u>) and the other targeting frontocentral theta-beta rhythms (<u>Farnia et al., 2015</u>), which suggests that some protocols may be effective, although the overall evidence remains inconclusive.

Across brain regions, NFT targeting the frontal cortex produced moderate effects, while parietal targets yielded small effects, and central or occipital targets showed no significant impact. The prefrontal cortex is involved in working-memory maintenance and strategic retrieval—particularly for verbal material (<u>Smith et al., 1998</u>)—and exerts top-down control that may facilitate hippocampal encoding, potentially explaining the observed STM advantages (<u>Simons & Spiers, 2003</u>). Parietal cortices contribute to information integration and are components of frontoparietal control networks implicated in attention and short-term memory, which may account for smaller but detectable effects at parietal targets (<u>Ptak,</u> <u>2012</u>).

For EEG-based NFT, alpha-band targeting (8–12 Hz) was associated with modest, consistent enhancements in memory performance, whereas theta (4–8 Hz) and beta (13–30 Hz) did not show significant effects when analyzed by frequency band alone. This pattern aligns with reports linking alpha oscillations to attentional modulation and efficient encoding (<u>Klimesch, 1999</u>; <u>Bazanova & Vernon, 2014</u>). Alpha upregulation may support STM and LTM by modulating encoding and enhancing selective and sustained attention (<u>Sitaram et al., 2017</u>). Alpha activity may also coordinate distributed cortical networks, thereby facilitating memory consolidation and working memory retention (<u>Benna & Fusi, 2016</u>; <u>Roux &</u> <u>Uhlhaas, 2014</u>). Although mechanistic interpretations remain tentative, the current body of evidence supports alpha modulation as a promising target for NFT-based memory enhancement. However, because subgroup analyses treat each factor independently, they may miss the co-occurrence structure that is captured by MCA (see Section 4.6).

### 4.4 Modality differences: EEG vs. fNIRS

EEG-based NFT showed small yet statistically significant benefits for memory, whereas fNIRS-based NFT did not. A major contributing factor to this discrepancy is likely the substantial difference in available evidence. Of the 24 studies included in the meta-analysis, EEG-based NFT was employed in 21 (*n* = 542), whereas fNIRS was used in only 3 (*n* = 107). Thus, although the current results favor EEG-based NFT for memory enhancement, this conclusion should be interpreted with caution. More well-powered, rigorously designed fNIRS studies are needed to allow for a definitive comparison between modalities.

Beyond sample size limitations, methodological differences between EEG and fNIRS may further account for the variation in observed outcomes. EEG provides millisecond-level temporal resolution, making it ideally suited to track and modulate fast neural oscillations (e.g., theta and alpha rhythms) implicated in memory encoding and attentional regulation (<u>Riddle et al., 2020</u>). In contrast, fNIRS measures slow hemodynamic responses, which may not align closely with the rapid neural activity required for oscillation-targeted NFT.

### 4.5 Meta-regression analyses

The results indicated a positive association between daily NFT duration and effect size, supporting the presence of a dose–response relationship. Importantly, the lower bound of the 95% confidence interval for the estimated effect exceeded 0.2 at 30 minutes per session—a commonly cited threshold for a small but meaningful effect (<u>Cohen, 1988</u>). This finding may inform future trial design by identifying a practical minimum for session length. Whether longer sessions consistently yield stronger cognitive outcomes remains to be determined.

We also tested whether NFT-induced gains diminish as a function of time since the last training session. The analysis did not identify a significant association between effect size and the timing of post-intervention memory assessments. This null result should be interpreted with caution, as most studies conducted outcome assessments either immediately after or within 24 hours of the final NFT session, and few included longer-term follow-up. Consequently, the available data were insufficient to determine whether NFT effects persist, decay, or necessitate booster sessions. Future longitudinal research is essential to clarify the temporal dynamics of NFT-induced memory enhancement.

### 4.6 Insights from the MCA factor map

The MCA revealed several characteristic cross-category associations. The first notable pattern was the proximity of frontal-theta to the “Effective” category point. In this sample, all theta-targeted neurofeedback studies employed frontal EEG electrodes and evaluated verbal memory, with four out of five reporting statistically significant improvements (<u>Enriquez-Geppert et al., 2014</u>; <u>Eschmann et al., 2020</u>; <u>Farnia et al., 2017</u>; <u>Rozengurt et al., 2017</u>). This clustering aligns with prior evidence linking prefrontal theta activity to executive control and working memory (<u>Cavanagh & Frank, 2014</u>; <u>Mitchell et al., 2008</u>). Second, parietal-alpha formed another cluster, consistent with literature associating posterior alpha rhythms with visual processing, attentional modulation, and large-scale network coordination (<u>Klimesch,</u> <u>1999</u>). Although parietal-alpha was positioned farther from the “Effective” category than frontal-theta, posterior alpha neurofeedback may still confer cognitive benefits and merits further targeted investigation. Third, central-beta appeared closer to the visuospatial domain and the “Null” category point, suggesting limited efficacy of beta-band training for visuospatial outcomes. MCA captures co-occurrence patterns among study features that may not map one-to-one onto single-factor subgroup effects, which helps explain why a frontal-theta cluster can coexist with an alpha-band effect in subgroup analysis.

Age and task-related patterns on the factor map mirrored the subgroup meta-analysis: younger adults were located closer to the “Effective” category point than older adults, and verbal/STM outcomes clustered nearer to “Effective” than visuospatial/LTM outcomes. However, MCA is inherently exploratory and descriptive, and its spatial geometry can be sensitive to sparsely populated categories. Given the modest number of contributing studies, these proximities should be interpreted with caution and regarded as hypothesis-generating patterns for targeted confirmation in future trials.

### 4.7 Limitations

Several methodological limitations of this meta-analysis and broader gaps in the NFT literature merit attention. First, only one of the included studies was preregistered, which raises concerns about selective reporting bias and the replicability of findings. Many studies lacked double-blind designs, increasing the risk that observed effects were inflated by expectancy or experimental bias. Future trials should adopt rigorous experimental designs—specifically randomized, double-blind, sham-controlled protocols with preregistered analytic plans.

Second, short intervention durations and limited follow-up windows constrain the interpretability of memory improvements. Most studies assessed outcomes within 24 hours of training. In the absence of longer-term follow-up, it remains ambiguous whether observed benefits are transient, cumulative, or require maintenance. Longitudinal study designs are essential for determining the durability, decay trajectory, or need for booster sessions of NFT-induced cognitive effects.

Third, protocol heterogeneity remains a substantial challenge. NFT designs varied widely in session length (3 to 90 minutes) and total duration (1 day to 18 weeks), making it difficult to identify optimal training parameters. While our meta-regression suggested that sessions lasting 30 minutes or more may be associated with stronger effects, this finding was only marginally significant and requires replication in larger, well-powered studies.

Fourth, subgroup analyses suggested diminished NFT efficacy in older adult cohorts. This observation must be interpreted with caution, as it may stem from multiple confounding factors. These include small sample sizes (<u>Button et al., 2013</u>) and greater cognitive heterogeneity in neuroplasticity within older cohorts compared to young adults (<u>Park & Bischof, 2013</u>; <u>Coutrot et al., 2018</u>). Future research should explore whether age-specific adaptations to NFT protocols may enhance responsiveness in older populations.

Finally, most studies included in this review used EEG-derived power spectra (e.g., alpha or theta amplitude) as feedback features. Event-related potential (ERP)-based NFT and slow cortical potential protocols were absent—likely due to limitations in our search strategy—even though they have shown efficacy in attention modulation and clinical applications (Doehnert et al., 2008; Sitaram et al., 2017). These protocols, particularly those targeting ERP markers for attention or working memory such as the P300 or contralateral delay activity (<u>Polich & Kok, 1995</u>; <u>Vogel & Machizawa, 2004</u>), may hold untapped potential for memory enhancement.

## 5. Conclusions

This systematic review and meta-analysis suggest that NFT modestly enhances memory performance in healthy adults and is associated with a favorable safety profile. However, the limited reporting of adverse events precludes definitive conclusions regarding the safety of EEG- and fNIRS-based NFT protocols. Effects were more pronounced under specific conditions, particularly in younger participants, for EEG-based protocols, on STM and verbal memory, and with sessions lasting at least 30 minutes.

These findings underscore the need for more rigorous and standardized experimental designs. Critical methodological priorities include preregistration, implementation of double-blind protocols, and inclusion of long-term follow-up assessments to assess the persistence of NFT-related memory improvements. Additionally, the MCA results highlight frontal theta upregulation as a particularly promising neurofeedback target for memory enhancement.

The advancement of next-generation neurofeedback should move beyond single-band and single-site paradigms. Multiband, network-level, and individualized NFT strategies offer considerable promise, particularly when integrated with emerging technologies such as machine learning, multimodal neuroimaging, and portable neurotechnology (<u>Mushtaq et al., 2024</u>). This study provides not only a quantitative synthesis of the current literature but also a conceptual foundation for guiding the design of more effective and scientifically grounded NFT interventions.

## Author contributions

Makoto Hagihara: Conceptualization, Data curation, Formal analysis, Investigation, Methodology, Software, Validation, Visualization, Writing – original draft, Writing – review & editing.

Reiji Ohkuma: Conceptualization, Data curation, Formal analysis, Investigation, Methodology, Software, Validation, Visualization, Writing – original draft, Writing – review & editing.

Tatsuro Fujimaki: Data curation, Investigation, Validation.

Masako Tamaki: Conceptualization, Investigation, Supervision, Validation.

Mitsuaki Takemi: Funding acquisition, Investigation, Project administration, Supervision, Writing – original draft, Writing - review & editing.

Maro G. Machizawa: Conceptualization, Data curation, Formal analysis, Methodology, Investigation, Project administration, Supervision, Validation, Writing – original draft, Writing - review & editing.

## Funding

This work was conducted as a part of Braintech Guidebook development in JST Moonshot R&D to Mitsuaki Takemi (Grant Number JPMJMS2012).

## Competing interests

Maro G. Machizawa is a founder and CEO of Xiberlinc Inc., and he receives a salary from and holds shares in Xiberlinc Inc. The other authors declare no competing interests.

## Data availability statements

Data and codes used in this study were shared in OSF (https://osf.io/9zpa6/).

## Acknowledgments

The authors thank the members of the Evidence Evaluation Committee for Braintech Guidebook, a specially organized group for the Moonshot R&D project, for their comments on the manuscript.

